# Origin and diversity of *Capsella bursa-pastoris* from the genomic point view

**DOI:** 10.1101/2023.07.13.548917

**Authors:** Aleksey A. Penin, Artem S. Kasianov, Anna V. Klepikova, Denis O. Omelchenko, Maksim S. Makarenko, Maria D. Logacheva

## Abstract

**Background:** *Capsella bursa-pastoris*, a cosmopolitan weed of hybrid origin, is an emerging model object for the study of early consequences of polyploidy, being a fast growing annual and a close relative of *Arabidopsis thaliana*. The development of this model is hampered by the absence of a reference genome sequence.

**Results:** we present here a subgenome-resolved chromosome-scale assembly and a genetic map of the genome of *Capsella bursa-pastoris*. It shows that the subgenomes are mostly colinear, with no massive deletions, insertions or rearrangements in any of them. A subgenome-aware annotation reveals the lack of genome dominance – both subgenomes carry similar number of genes. While most chromosomes can be unambiguously recognized as derived from either paternal or maternal parent, we also found homeologous exchange between two chromosomes. It led to an emergence of two hybrid chromosomes; this event is shared between distant populations of *C. bursa-pastoris*. The whole-genome analysis of 119 samples belonging to *C. bursa-pastoris* and its parental species *C. grandiflora/rubella* and *C. orientalis* reveals introgression from *C. orientalis* but not from *C. grandiflora/rubella*.

**Conclusions:** *C. bursa-pastoris* do not show genome dominance. In the earliest stages of evolution of this species a homeologous exchange occurred; its presence in all present-day populations of *C. bursa-pastoris* indicates on a single origin of this species. The evidence coming from whole-genome analysis challenges the current view that *C. grandiflora/rubella* was a direct progenitor of *C. bursa-pastoris*; we hypothesize that it was an extinct (or undiscovered) species sister to *C. grandiflora/rubella*.

## Background

Polyploidization, or whole genome duplication, is a recurrent trend in the evolution of eukaryotic genomes (Van de Peer *et al*., 2017). It is especially prevalent in plants – in this lineage it greatly contributed to the morphological diversity and ecological adaptations (Madlung, 2013). Polyploidy can also influence such critical traits as mating system (self-incompatible vs self-compatible), tolerance to abiotic stresses and growth rates. It was also shown that polyploidy is a key factor of plant domestication (Salman-Minkov *et al*., 2016). It is thus thoroughly studied, however many important points concerning polyploids are not well understood. While the main model object of plant genetics, *Arabidopsis thaliana*, is not a polyploid, much attention has focused on the relatives of *A. thaliana* that display auto- or allopolyploidy (Burns *et al*., 2021; Monnahan *et al*., 2019; Han *et al*., 2015). Their close relationships to *A. thaliana* enhances the characterization of genomes and the application of modern functional genetics tools such as gene silencing and genetic modification. Among these species, the most intriguing is *Capsella bursa-pastoris*. This is an allopolyploid; after a long discussion about its origin its parental species were found to be *Capsella rubella/grandiflora* (these two species diverged very recently, many years after the emergence of *C. bursa-pastoris*) and *Capsella orientalis* (Douglas *et al*., 2015), two species that diverged about 1 Mya (Douglas *et al*., 2015). Each parent has relatively small habitat regions - *C. grandiflora* is confined to mediterranean region (https://powo.science.kew.org/taxon/urn:lsid:ipni.org:names:279976-1 Accessed 26 June 2023) and *C. orientalis* - to Eastern and Sourthern Russia, Kazakhstan, Mongolia and China (https://powo.science.kew.org/taxon/urn:lsid:ipni.org:names:280028-1 Accessed 26 June 2023). Surprisingly, their hybrid is a cosmopolitan weed growing around the globe – from Arctica on the North and Kerguelen islands on the South (Frenot *et al*., 2005) (https://powo.science.kew.org/taxon/urn:lsid:ipni.org:names:30092589-2 Accessed 26 June 2023). It is one of the most widespread plants on Earth, the ecological niches that it inhabits range from African deserts to polar lands. Such plasticity has been at the focus of the studies in ecology and ecological genetics (Choi *et al*., 2019; Wesse *et al*., 2020). It is presumably mediated by polyploidy; earlier some genetic cues that might be involved in environmental adaptation were found, they include differential splicing and asymmetry of regulatory elements between subgenomes (Kasianov *et al*., 2017; Slotte *et al*., 2009) though this aspect is very far from being well understood. *C. bursa-pastoris* also has considerable morphological variation, including the floral phenotypes that are not found in *A. thaliana* and rarely – in other Brassicaceae [12, 13], thus being promising object for the study of development (Hintz *et al*., 2006). The main obstacle to using *Capsella bursa-pastoris* as a model object was is lack of a well-assembled genome. Despite the significant progress in sequencing there is still no chromosome-scale assembly of *C. bursa-pastoris* genome; most analyses are done using mapping on a reference genome of *C. rubella*, one of the parental species, and further phasing into subgenomes (Kryvokhyzha, Milesi, *et al*., 2019). This leads to biases due to the unequal efficiency of mapping of the reads belonging to the O (the one derived from *C. orientalis* parent) and R (derived from *C. rubella*/*grandiflora*) subgenomes.

This paper presents the assembly of the *Capsella bursa-pastoris* genome down to the chromosome level, obtained by a combination of HiFi Pacbio reads, HiC, and high-resolution genetic mapping. The genome-wide analysis showed that this species has single origin without active introgression of genes from parental species and admixture between geographically isolated accessions.

## Results

Our approach to the construction and correction of the assembly is a multistep procedure that includes the integration of chromosome conformation capture (HiC) data and the genetic map (Figure 1). At first step we obtained contigs from Pacbio CCS reads. Total length of this initial assembly was 363 941 kb with N50 ∼ 1 251 kb. Since the subgenomes have very high sequence similarity we may expect the chimeric contigs combining regions from homeologous chromosomes. Thus we performed an assembly check using the data on segregation of markers in F2 populations derived from the cross of two polymorphic accessions – lel and wt3.4-msk (Supplementary figure 1) (Kasianov *et al*., 2017; Klepikova *et al*., 2021). These data, obtained using WGS, contained ∼ 254 thousands of SNP markers with average distance between markers 864 bp. We analyzed the character states for these markers in 50 plants from F2 population (see example on Supplementary figure 2). If more than two recombination was observed between two adjacent markers, the sequence containing these markers was treated as misassembly (see example on Supplementary figure 3). The contig/scaffold was then split and the sequence between markers was removed from the assembly. We found 71 such cases; in most of the cases the number of observed (false) recombinations was more than 25% - that supports the markers are in fact not linked and are inherited independently. As a result N50 reduced to 1 063 kb. Then we proceeded to scaffolding using HiC data. The HiC reads are short and frequently thus cannot be unambiguously assigned to one of the subgenomes for many genome regions. In order to overcome this limitation we again used genetic map data and assigned contigs to linkage groups. 490 contigs were assigned to 16 linkage groups corresponding to 16 chromosomes typical for *C. bursa-pastoris*. N50 of contigs assigned to linkage groups was 1 312 kb and total length of the assembly ∼ 319 Mb. After that we performed scaffolding on each linkage group separately. Most contigs were scaffolded, for only four short contigs together making 0.5 Mb the scaffolding was not successful. After this we mapped HiC data on the chromosome scaffolds and performed visual examination and check of the assembly quality. Based on this check we removed to the regions which showed irregular or ambiguous position based on HiC map (Supplementary figure 4). These regions mostly correspond to the repeat-rich fraction of the genome. At this step we removed 66 Mb from the assembly. The resulting assembly consisted of 16 chromosomes with N50 16.6 Mb and total length ∼ 253 Mb. This is less than the size estimated based on cytofluorimetry (Lysak *et al*., 2009), however, as we show below, it contains the overwhelming majority of genes.

**Figure 1.**
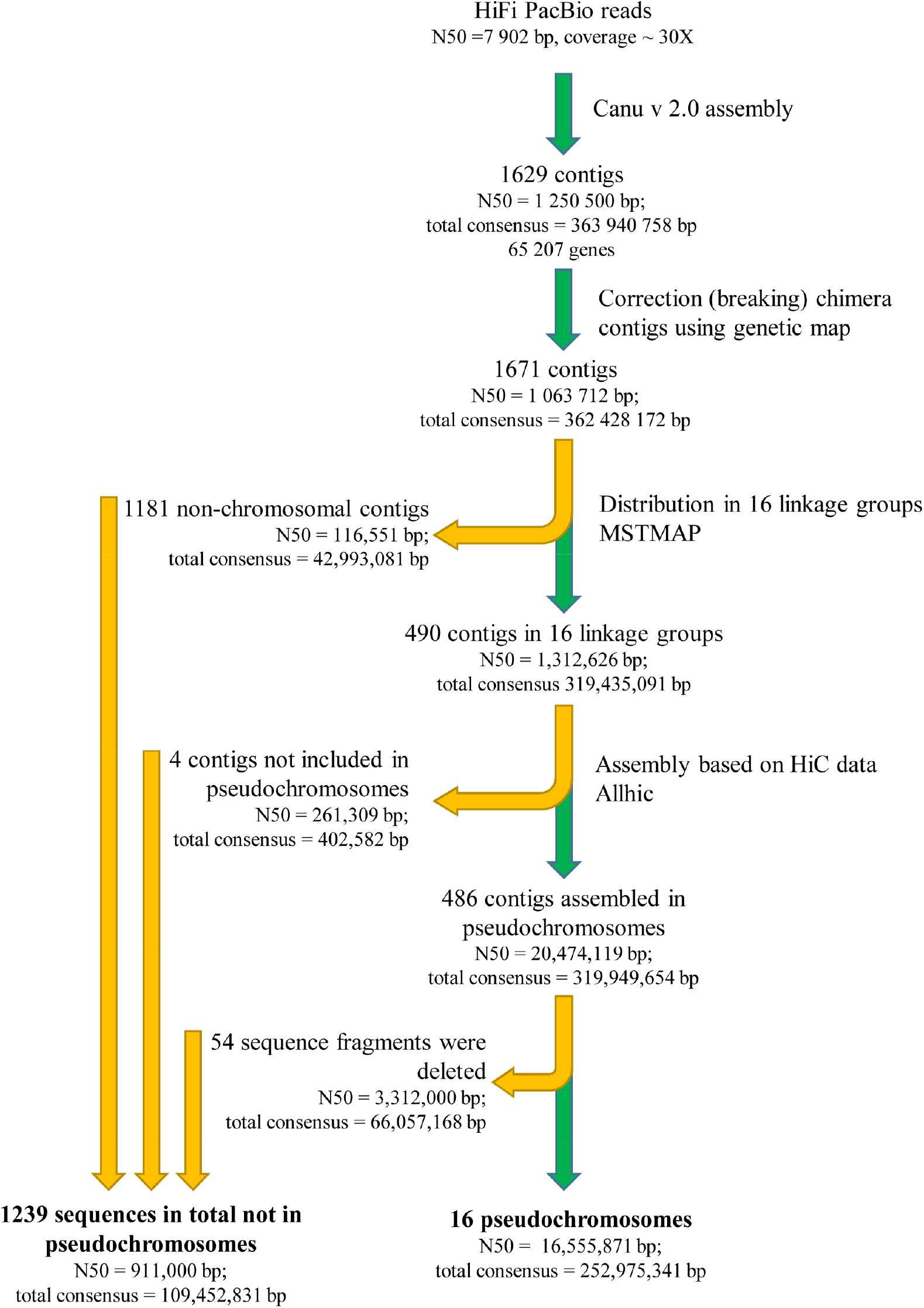
Assembly pipeline for the *Capsella bursa-pastoris* genome.

At the next step we assigned chromosomes to subgenomes. To do this we applied approach based on the mapping of the reads from parental species *Capsella rubella* and *Capsella orientalis*. In order to assess its validity we performed a simulation of *C. bursa-pastoris* genome using the sequences of parental genomes: we combined sequences of *C. rubella* and *C. orientalis* genomes into single reference and then mapped the reads of each of the species. The analysis of mapping rates showed that most part of the reads from each species is mapped on the subgenome where is belongs to, not on the subgenome from other parent, despite their high similarity (Supplementary figure 5). This allows to use mapping rate as a proxy for the determination of the origin of chromosomes from either R or O parent. Thus we mapped the reads of *C. rubella* and *C. orientalis* on *С. bursa-pastoris* chromosomes and calculated coverage on 10-Kb windows (Figure 2a). This showed that most of the chromosomes are derived from one parent exclusively (i.e. there were no recombination between R and O subgenome after the formation of a hybrid) with exception of two chromosomes that carry an evidence of homeologous exchange between R and O subgenomes chromosomes (these two chromosomes are termed here and further O7_R7 and R7_O7). The genetic map also supports the hybrid nature of these chromosomes. After obtaining corrected assembly we annotated it using a combination of homology based and de novo prediction approaches. The annotation included 65,207 protein-coding genes, 637,76 (97.8 %) out of which are located in the chromosome scaffolds and are evenly distributed across subgenomes (31,875 in the subgenome O and 31,901 in the subgenome R). In order to facilitate further studies employing this annotation the genes were put into correspondence with orthologous *Arabidopsis thaliana* genes. The gene naming reflects this correspondence following the format: (number of chromosome).(number of gene in *Capsella*).(number of gene in *A. thaliana*)_(type of search of a homologous gene - orthofinder or blast), for example, «Cbp.O1.g00001000.AT1G02140_O» for a gene that is located in the chromosome O1, is a first gene annotated on this chromosome and is orthologous to *A. thaliana* gene AT1G02140. The genes are numbered with space of 1000 in order to enable the addition of genes that might be found in other accessions of *С. bursa-pastoris* in further studies.

In order to perform the quality control of the assembly in terms of completeness of gene set we estimated the occurrence of all annotated genes and the genes from BUSCO set in different fractions of the assembly: the initial assembly, the subgenomes of the final assembly and the sequences that were deleted at different steps of the curation (for example, the misassembled regions and repeat-rich regions). This analysis demonstrated that fraction of contigs not included in the final (pseudochromosomes) assembly contained less than 5% of BUSCO genes and less than 2% of all genes (Figure 2b, Supplementary table 1). This means that the fragments not included in the final assembly correspond to the gene-poor fraction of the genome (presumably pericentromeric regions) and do not affect further analysis unless they are focused on repetitive elements. Another approach to quality control, aimed at the inference of the correctness of the assembly is based on the HiC data. Visual exploration of the contact map shows that it has typical diagonal patterns with no regions that may indicate on misassembly (Figure 2c, Supplementary figure 2).

**Figure 2.**
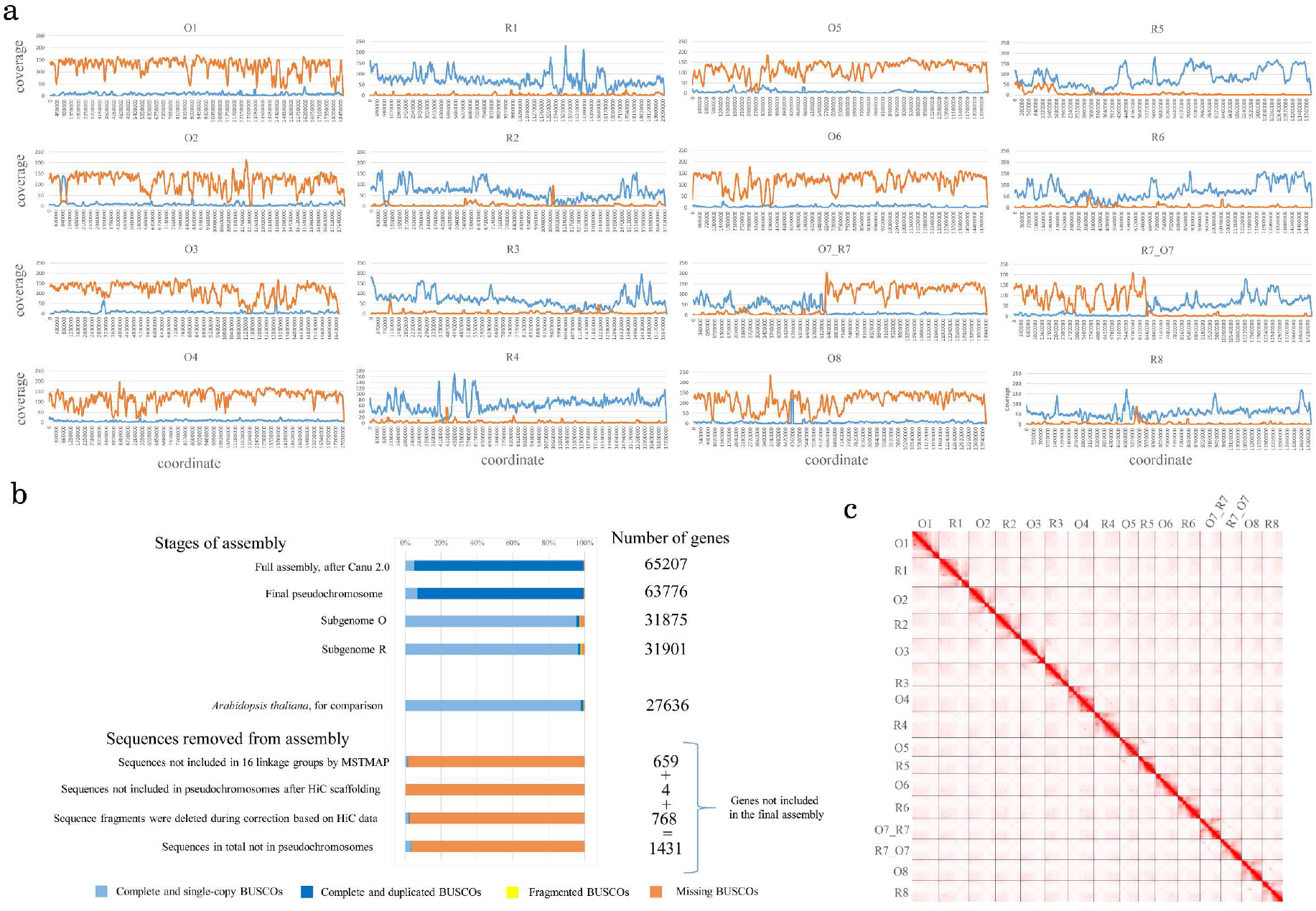
*Capsella bursa-pastoris* genome assembly. a) Coverage of *С. bursa-pastoris* chromosomes by the reads of *C. orientalis* (orange) and *C. rubella* (blue) b) Analysis of the representation of BUSCO genes and all protein-coding genes in chromosomes and fragments of genome not included in chromosomes. c) HiC map of *C. bursa-pastoris* chromosomes.

The R and O subgenomes are mostly colinear one with another and with the *C. rubella* parent (the only one for which chromosome-scale assembly is available). As whole genome alignments show, there is no major rearrangements or INDELS between homeologous chromosomes (Figure 3a). Many polyploids of hybrid origin have undergone biased fractionation of the subgenomes, where one of the subgenomes preferentially loses genetic material and other preferentially retains. This is clearly not the case for *C. bursa-pastoris*. This finding is congruent with the results stemming from the annotation: both subgenomes carry similar number of genes. The distribution of genes and repeats is also similar; the repeats have a sharp peak at a specific gene-poor region that presumably corresponds to a centromere. The only notable difference is the small fragment at the beginning of chromosome R5 for which there is no homologous fragment in O subgenome (neither in O5 not in any other place of the subgenome). In order to ensure that this is not an assembly artifact we checked its coverage in R5 and it does not deviate from the average across the genome; we also looked for its presence in the contigs of the initial assembly that were not included in the final one and did not found any. This region is found in *C. rubella* genome; this suggests that the absence in O subgenome is the result of deletion occurred after the divergence *C. grandiflora/rubella* and *C. orientalis*. The *C. rubella* genome has a small inversion in the beginning of chromosome 4 relative to both R and O subgenomes (Figure 3b). The fact that R and O subgenomes share the same orientation shows that it was ancestral while the inverted orientation found in *C. rubella* is lineage-specific and occurred during speciation of *C. rubella* after the formation of *C. bursa-pastoris*.

**Figure 3.**
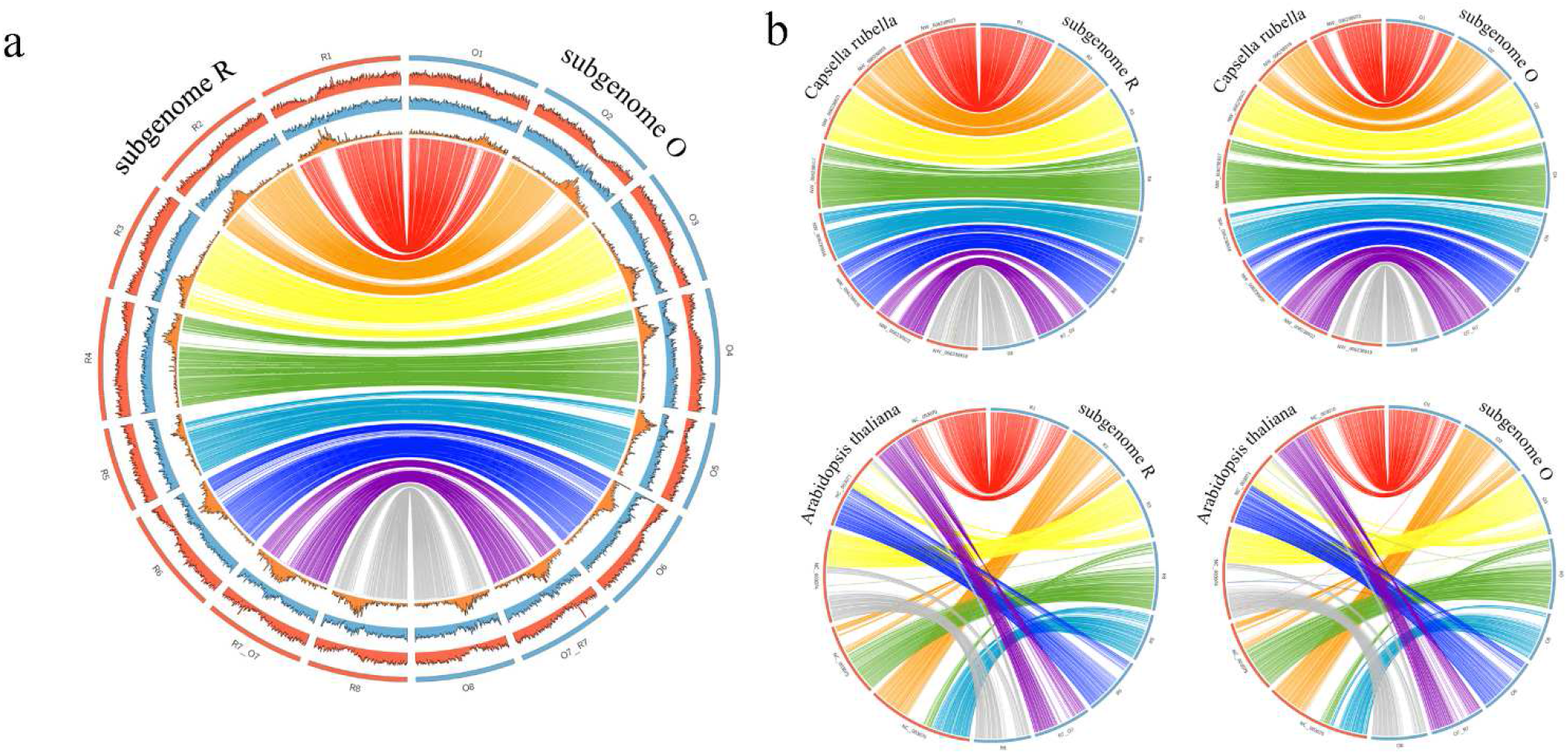
Circos plots showing the comparison of *С. bursa-pastoris* subgenomes. a) between themselves b) with *С. rubella* and *A. thaliana*. On the panel a, the density of repeats is shown in orange, the density of genes in blue and GC content in red.

Both *C. bursa-pastoris* subgenomes retain strong similarity and colinearity with *A. thaliana* genome (Figure 3b), with *A. thaliana* chromosomes aligning to several chromosomes of each of the subgenomes. This is in congruence with previous data on *C. rubella* (Anon, n.d.; JOHNSTON *et al*., 2005) and highlights two contrasting tendencies in the genome evolution in Brassicacae – whole genome duplication and fusions of chromosomes with further reduction of non-coding DNA.

The availability of a subgenome-resolved genome sequence and a wide set of WGS data on *C. bursa-pastoris* and parental species derived from earlier studies (Huang *et al*., 2018; Kryvokhyzha, Salcedo, *et al*., 2019) allows to analyze the evolution of each of the subgenomes separately. To do this we used the samples from species that is donor of the other subgenome as outgroup. For example, when considering O subgenomes, *C. rubella* and *C. grandiflora* were used as outgroup and vice versa. The reads of diploid species were mapped on only one subgenome – the one that was under consideration. For *C. bursa-pastoris* samples we mapped reads on both subgenomes and then considered only the regions corresponding to one subgenome. Using this approach we got 6 651 155 variable positions (SNP, INDEL were not used) for subgenome R and 4 967 014 for subgenome O on a set of 56 samples of *C. bursa-pastoris*, 7 samples of *C. rubella* and 21 for *C. grandiflora*, and 15 for *C. orientalis* (Additional file 7, Figure 4d). These genome-wide data on genetic variation were used to infer relationships between samples using phylogenetic trees and neighbor-net approach. All these analyses were carried out separately for R and O subgenomes. In terms of the relationships within *C. bursa-pastoris*, phylogenetic trees resulting from the analysis of O and R subgenomes are mostly congruent. Both show the separation of Asian (ASI) and Middle East (ME) populations recognized in earlier studies (Huang *et al*., 2018; Kryvokhyzha, Salcedo, *et al*., 2019); the third group, European (EU) is recognized as monophyletic only in trees inferred from O subgenomes while in R-trees it is present as a grade while ASI and ME are sister clades (Figure 4a). Neighbor-net analysis offers higher resolution and allows to get more detailed and accurate view of the relationships between *C. bursa-pastoris* populations. The networks inferred from different subgenomes are highly congruent (Figure 4a) and show that *C. bursa-pastoris* is subdivided into six subgroups. Three of them correspond to the EU + ME populations and are found in Europe, with group 1 confined mostly to northern Europe and groups 2 and 3 – to Eastern (the group 3 corresponds to ME). Other three occur in Asia (group 4 – northern Kazakhstan and groups 5 and 6 – China). A notable feature of both networks and trees is the asymmetry concerning the position of *C. orientalis* relative to the subgenome O and *C. rubella*/*C grandiflora* to the subgenome R. For the R-lineage (with *C. orientalis* is a root) *C. rubella*/*C grandiflora* is an outgroup to R subgenomes. In case of O-lineage (*C. orientalis* and the subgenome O, *C. rubella*/*C grandiflora* is a root) it is different – the clade uniting *C. orientalis* samples is nested together with the subgroups of *C. bursa-pastoris*. Similar observations were done in earlier studies (Kryvokhyzha, Milesi, *et al*., 2019) and two hypotheses were put up: the multiple origin of *C. bursa-pastoris*, independent origin of Asian *C. bursa-pastoris* and a present-day *C. orientalis* being closer to the ancestor of Asian group and the introgression from *C. orientalis* to the O subgenome. The availability of a genome sequence phased into subgenomes allows us to test those hypotheses and to put up an alternative one (see discussion). As mentioned above, we found a homeologous exchange between chromosome 7 of *C. orientalis* origin and that of *C. rubella/grandiflora* origin. It led to a formation of two hybrid chromosomes, termed R7_O7 and O7_R7. The accession that we use as a source for reference (msk3-4) belongs to the EU population (Kryvokhyzha *et al*., 2016) or “group 1” according to the classification presented above. To test whether the hybrid chromosomes are found in Asian and Middle East groups or it is a feature unique for this accession we performed the analysis based on chromatin contacts data acquired using HiC method (Lieberman-Aiden *et al*., 2009). We constructed and sequenced HiC libraries for representatives of Asian (Cbp_ASI) and Middle East (Cbp_ME) groups and mapped them on msk3-4 reference. For both Cbp_ME and Cbp_ASI contact maps for R7_O7 and O7_R7 demonstrate a clear diagonal pattern with no gaps. This indicates on the absence of structural changes between the reference genome and the genome from which HiC data are derived (Figure 5a). In order to control whether this analysis is able to reveal such changes we made a simulation of translocation (which will correspond to the absence of exchange between R7 and O7) by artificially modifying the reference and mapped HiC data to this modified reference. In this case the contact map clearly shows the gap in contacts which is an indicator of the discordance between the reference and the HiC data (Figure 5a). As an additional evidence of the colinearity of genomes from different populations of *C. bursa-pastoris* and the presence of hybrid chromosomes we performed a crossing of accessions belonging to the populations 3 and 1 (the latter is the same accession msk3-4 that was used for the genome assembly). We analysed 30 F2 plants derived from two crosses and calculated the number of recombinations per chromosome (Figure 5b). In case if the chromosomes 7 had different structure we would have observed decreased recombination in these chromosomes. Taken together, these analyses provide the evidence of that hybrid chromosomes exist in all three major clades of *C. bursa-pastoris*.

**Figure 4.**
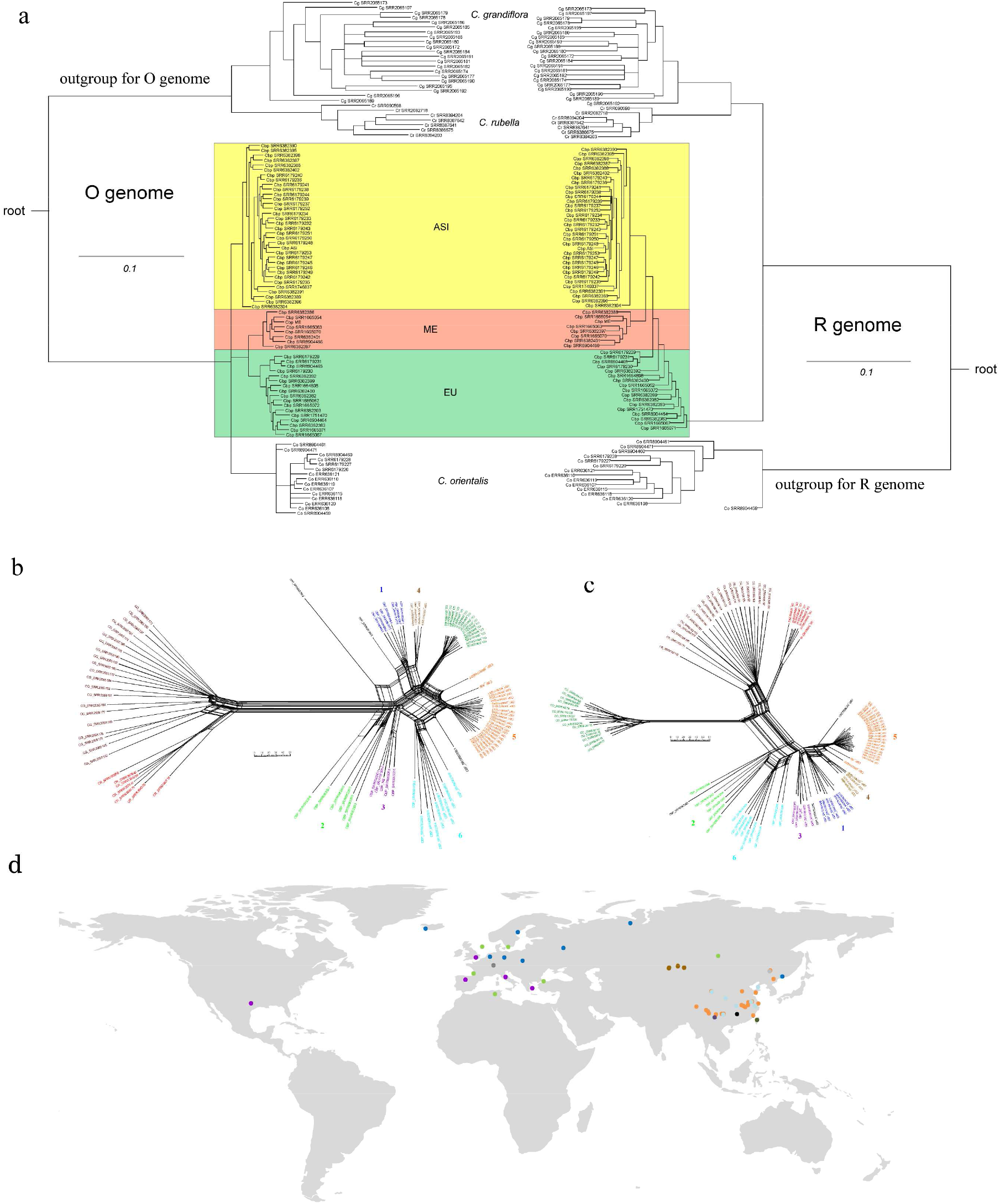
Populations of *C. bursa-pastoris.* a) tangelogram of phylogenetic trees by subgenomes O and R Colors indicate samples of groups ME, ASI, EU. b) neighbor-net graph constructed for subgenome O c) neighbor-net graph constructed of subgenome R d) geographical map with the location of populations. The six subgroups are each marked with their own color and number. Accessions belonging to the same groups on both trees are marked with the same color. Samples with an unstable position are highlighted in black. The color is the same on a geographic map and on graphs

**Figure 5.**
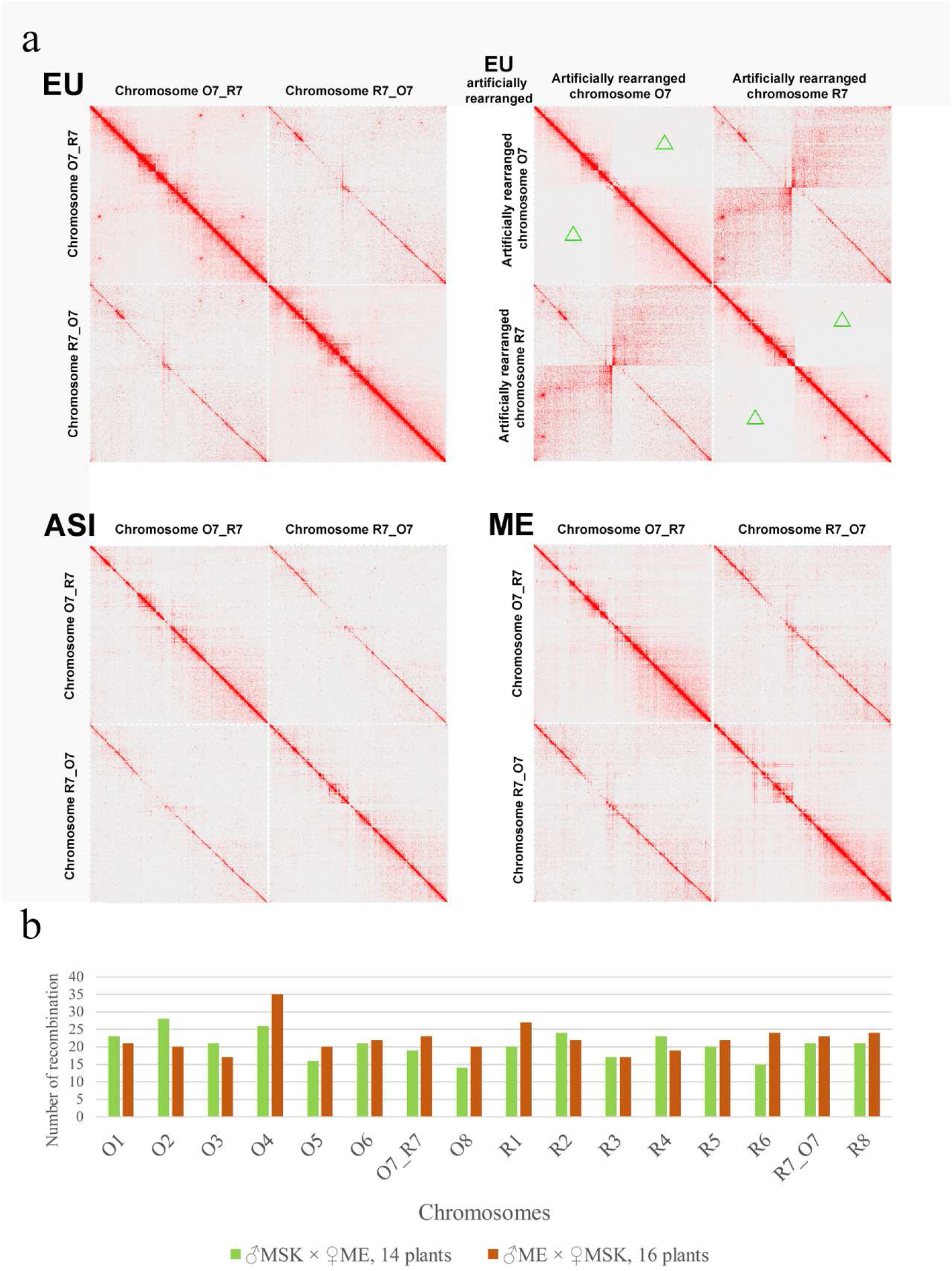
Homeologous exchange between chromosome 7 O and R subgenome. a) HiC maps for chromosome 7 for plants belonging to different populations (area without contacts in the analysis of the artificial chromosome is indicated by green triangles). Non-unique read mapping was allowed, in order to visualize the synteny of paralogous chromosomes. b) the number of recombinations in chromosomes detected in F2 from crossing line MSK (EU population) and line Cbp_ME (ME population).

Another important question is the transfer of genetic material between populations and from parental species. In order to assess it we analyzed population structure based on genome-wide search of SNPs. This was done separately for each of the subgenomes. Under k=4 (the optimal value as estimated by fastStructure) we did not found the evidences of widespread introgression from either parent to modern *C. bursa-pastoris* populations (Figure 6). Only the introgression of *C. orientalis* to O subgenome is observed in few samples from Asian populations (groups 4 and 5) which come from the areas overlapping with the areas of *C. orientalis* distribution (https://powo.science.kew.org/taxon/urn:lsid:ipni.org:names:280028-1 Accessed 26 June 2023). Previous reports also detected introgression though the details differ (will be discussed further). Surprisingly, the analysis of population structure revealed the signs of introgression from *C. bursa-pastoris* to parental species. This is highly unexpected because requires the hypothesis on differential elimination of one of the subgenomes in gametes; though such cases are known in some species (Dedukh *et al*., 2015) we think that this might also stem from technical issues (like contamination by *C. bursa-pastoris* material because these species are very hardly distinguishable) rather than from biological phenomena. Within *C. bursa-pastoris* the gene flow between populations was also moderate. Using admixture analysis under K = 6 (corresponds to the number of groups distinguished within *C. bursa-pastoris*) we found that R subgenomes (Supplementary figure 6) do not demonstrate any stratification while for O subgenomes two groups are separated (corresponding to EU + ME and Asian populations). Within these groups several samples show admixture with European populations. These samples correspond to the group 5 (Figure 4b) of the Asian clade. This might however be explained not only by introgression but also the retention of ancestral polymorphisms because this population represent the basal clade of the Asian group. In any case, there is no massive admixture between populations.

**Figure 6.**
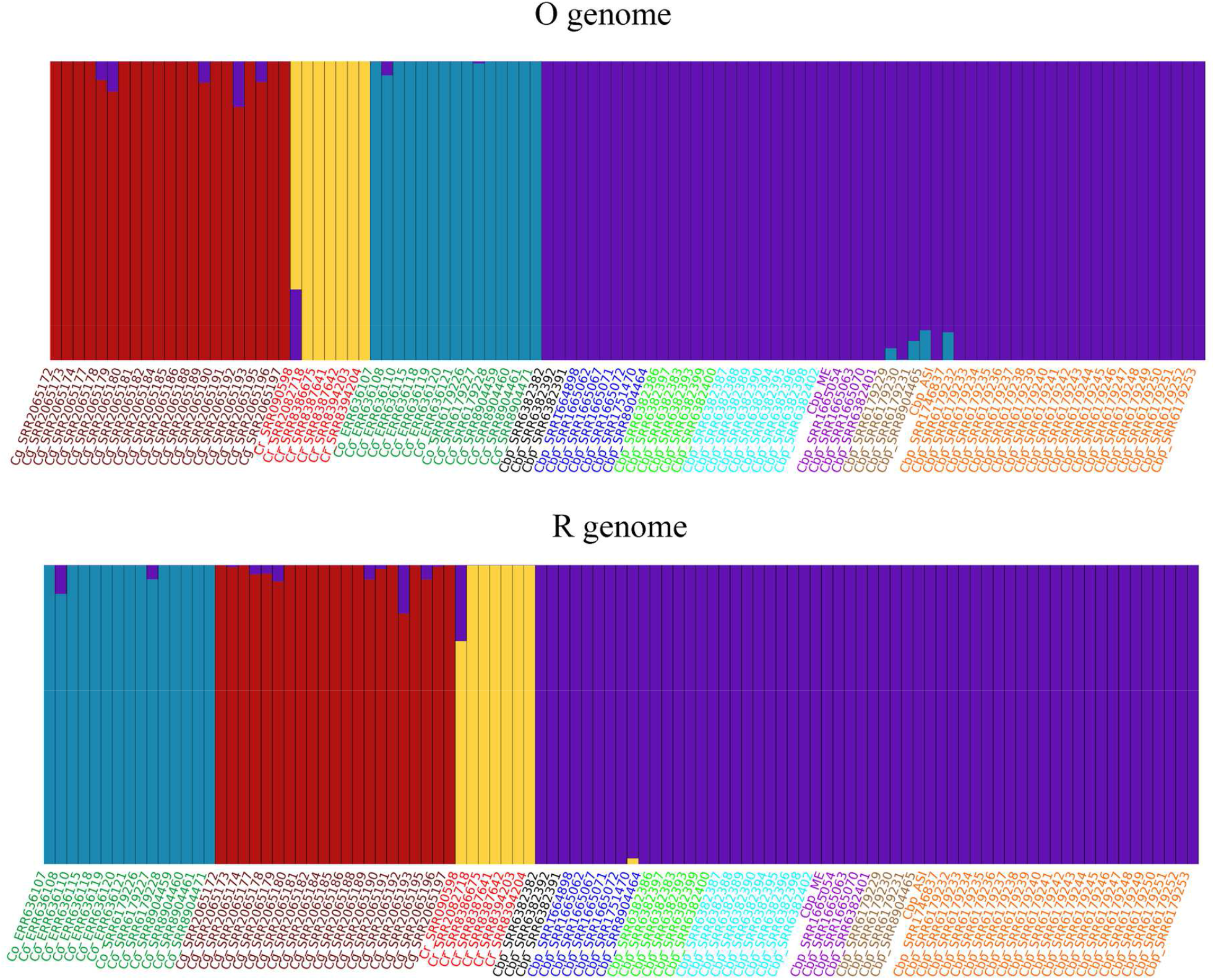
Analysis of introgression by admixture analysis, for K=4. The colors of the line names correspond to the populations in Figure 4b.

## Discussion

*C. bursa-pastoris* has long been a complex object. There was a long-standing controversy about its origin (either by allo- or by autopolyploidy (St Onge *et al*., 2012; Roux and Pannell, 2015; Hurka *et al*., 2012)) which was solved only when genome-scale data came into practice (Douglas *et al*., 2015; Kasianov *et al*., 2017). However many important questions concerning the evolution of this species remained. Some of these questions were tackled using a genome of one parental species - *C. rubella* - as a reference for phasing-based analysis (Kryvokhyzha, Salcedo, *et al*., 2019). Our analysis of chromosome-scale assembly allowed to characterize the structural features of each of the subgenomes. We found that they are mostly colinear and carry similar number of genes. In contrast to what is seen in many other polyploids of hybrid origin (Bird *et al*., 2018), there is no biased fractionation leading to the subgenomes dominance. This is congruent with our earlier findings on the absence of asymmetry in gene expression profiles between subgenomes (Kasianov *et al*., 2017).

An important finding from the whole genome analysis is the evidence of the homeologous exchanges (HE) between R and O subgenomes at early stages of *C. bursa-pastoris* evolution that led to a formation of two hybrid chromosomes. The homeologous exchanges are frequent in allopolyploids (Mason and Wendel, 2020) however the fixation of each individual exchange event is rare and the fact that HE event is shared is a strong evidence of common ancestry (Lashermes *et al*., 2014). The question on a single or multiple origin of *C. bursa-pastoris* is debatable; an independent origin was hypothesized, especially for Asian clade, though inconclusively (Kryvokhyzha, Salcedo, *et al*., 2019) We showed that the representatives of ME and Asian clade also have hybrid chromosomes, similar to msk3-4 accession belonging to European clade. Though we can not completely rule out the possibility that there are some accessions that lack HE or that the HE occurred independently in different clades, the current results strongly support the single origin of *C. bursa-pastoris*. Further study of *C. bursa-pastoris* accessions, at a pangenome level would corroborate this.

Notably, the phylogenetic trees and networks inferred from R and O subgenomes are discordant relative to the position of *C. orientalis* and *C. rubella*/*C grandiflora* samples. These distinct evolutionary trajectories could be the evidence of multiple origin (Kryvokhyzha, Salcedo, *et al*., 2019). However, in view of the presence of HE in different populations of *C. bursa-pastoris* this scenario is unlikely. It should also be noted that under the hypothesis on multiple origin the trees based on the topologies of O and R subgenomes are expected to be concordant (unless we hypothesize that *C. bursa-pastoris* originated from different populations of *C. orientalis* crossed with the same population of *C. rubella*/*C grandiflora*). Alternative explanation also put out in the earlier studies is the massive introgression from *C. orientalis* to *C. bursa-pastoris*. However we showed that this is not the case as well since the introgression is limited to the several samples from Asian populations. Taking into account our findings stemming from whole genome analysis we put out another hypothesis for this contrasting phylogenies. It implies that the *C. rubella*/*C grandiflora* is not direct progenitor (paternal parent) of *C. orientalis*, but a sister species that diverged from a species that is a direct progenitor and is now extinct or not identified (Figure 7). This is supported also by the higher divergence between Cr/Cg and R subgenome than between Co and O subgenome.

**Figure 7.**
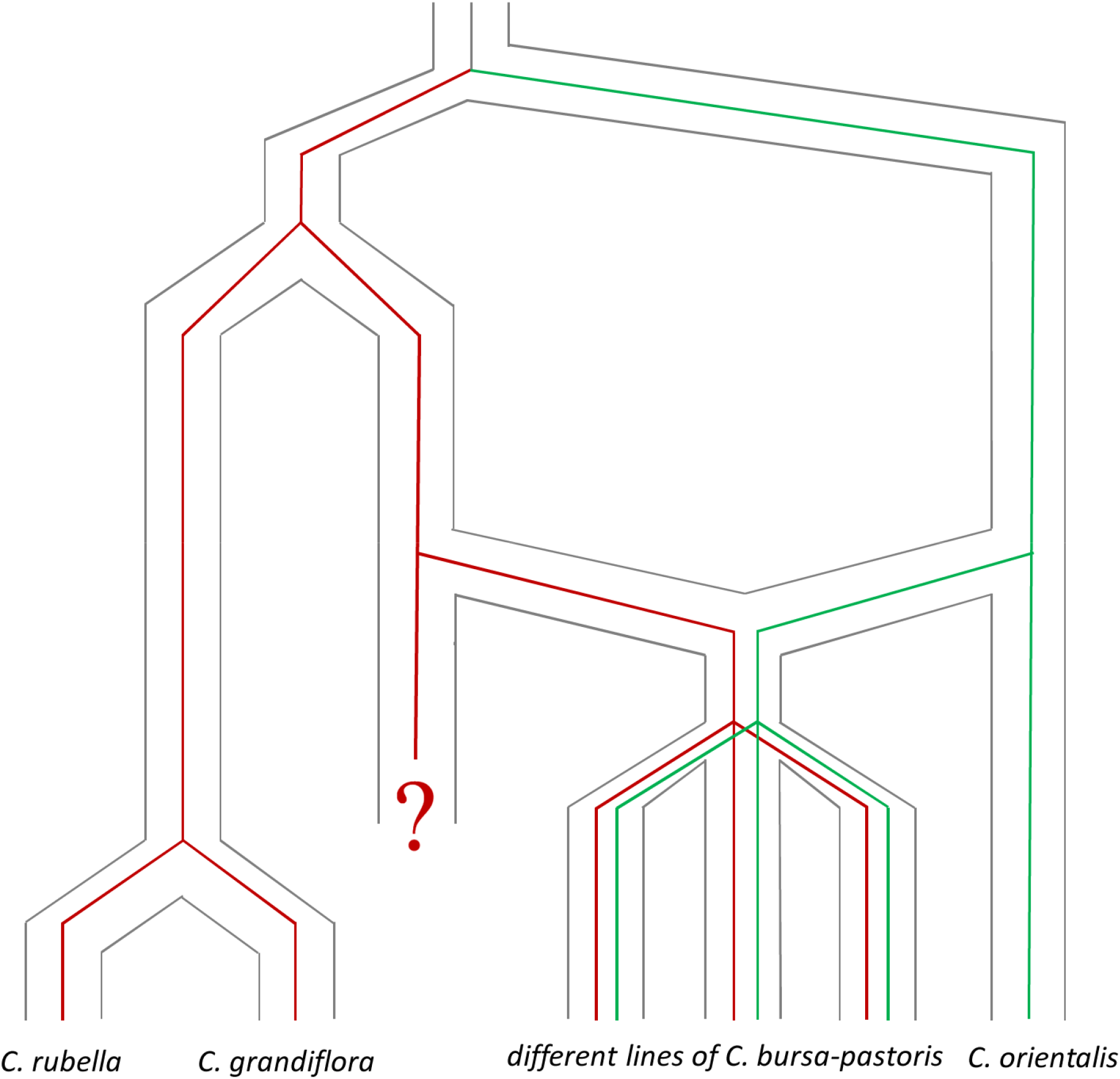
Evolution of the two subgenomes of the tetraploid *Capsella bursa-pastoris* (Brassicaceae).

The genome of *C. rubella* was by now the only one assembled up to chromosome scale; the genome of *C. orientalis* is available only as a set of short contigs (N50 = 25 Kb) (Ågren *et al*., 2014), and is not used for any studies involving *C. bursa-pastoris*. Instead of this, the multi-step pipeline involving mapping of *C. bursa-pastoris* reads on *C. rubella* genome and further phasing of SNP is used (Kryvokhyzha, Salcedo, *et al*., 2019). Given the higher divergence between *C. rubella* and R subgenome and the presumable non-parental relationships of *C. rubella/C grandiflora* and *C. bursa-pastoris* this might obfuscate the results of the genomic and transcriptomic analysis based on this pipeline. The availability of the chromosome-scale assembly of *C. bursa-pastoris* genome thus opens the avenue to the unbiased view of the genome evolution of this plant.

## Materials and methods

### Plant material, DNA extraction, library preparation, and sequencing

For PacBio long-read sequencing, fresh leaves of *C. bursa-pastoris* line ‘msu-wt’ (Klepikova *et al*., 2021) were quick-frozen in liquid nitrogen and transferred on dry ice to the DNA Link laboratory (South Korea, Seoul). DNA was isolated, and SMRTBell libraries were prepared and sequenced by the DNA Link laboratory. For WGS short-read sequencing which was performed for two accessions, termed here as Cbp_ASI and Cbp_ME, DNA was extracted from fresh leaves using CTAB method (Doyle and Doyle, n.d.). Shotgun libraries for genome assembly were prepared using the NEBNext Ultra II DNA library preparation kit (New England Biolabs, Ipswich, MA, USA) and sequenced on Illumina HiSeq2000 (Illumina, San Diego, CA, USA) platform using HiSeq SBS Kit (200 Cycles).

For chromatin capture-based Hi-C sequencing, the nuclei enrichment and isolation was performed on fresh leaf material using CelLytic PN Isolation/Extraction Kit (Sigma-Aldrich). DNA was extracted from plant cells and Hi-C libraries were prepared using EpiTect Hi-C Kit (Qiagen, Germany) according to the manufacturer’s protocol. Hi-C libraries were sequenced on Illumina NextSeq500 (Illumina, San Diego, CA, USA) platform using platform using High Output Kit v2.5 (150 Cycles).

The quantity of extracted DNA and prepared libraries were measured with Qubit (Thermo Fisher Scientific, Waltham, MA, USA) DNA assays. The size and quality of prepared libraries were validated by Bioanalyzer 2100 (Agilent, Santa Clara, CA, USA) DNA fragment analysis and qPCR.

### Genome assembly

Draft genome assembly of the *C. bursa-pastoris* genome was constructed from CCS PacBio reads (the average read length was 7,948 bp) obtained from the DNA Link laboratory using the Canu software v. 2.0 (Koren *et al*., 2017) with the following parameters: genomeSize=300m - estimated genome size; correctedErrorRate=0.005 - the allowed difference in the overlap between two reads is no more than 0.5%; minOverlapLength=1000 - minimum read overlap length.

The reads of 50 samples of the F2 generation and one of the parent plants of the F0 generation used for the genetic map were mapped to the assembly, and SNP calling using the CLC Genomics workbench software v.20.0.3 (QIAGEN) was performed to verify and correct the assembly (see Supplementary figure 7). Based on the SNP data of the F0 generation parent, the database of homozygous SNP markers (245,060 markers in total) was constructed and used for the downstream analysis (see Supplementary figure 8). The allele status within each of the F2 plants was tested based on SNP calling results and marker coverage information for each marker. The results of this check were compiled in tables where rows correspond to the genetic markers of the database based on F0 sequencing results, columns correspond to the F2 samples, and each table cell contains information about the marker state for the intersection. Data collection for the genetic map, SNP database construction, and data processing in detail is presented in Additional files 1,8,9. Marker states were as follows:

- “-1” – homozygous allele corresponding to the reference allele. The assembly from sequencing data of the first parent of the F0 plants of the genetic map was used as a reference.
- “0” – heterozygous
- “1” – homozygous allele corresponding to the homozygote from the second parent of the F0 plants of the genetic map. The marker database has been created from the data on the homozygotes of this plant.
- “-” – unidentified marker.

Before searching for incorrectly assembled regions, the table with the states of the genetic map markers was filtered. Markers that were unidentified in more than half of the samples were removed. Markers that did not satisfy Pearson’s criterion for the ratio of homozygotes to heterozygotes were also removed. The threshold for the criterion was chosen as 5.99, corresponding to a p-value < 0.05. Next, the filtered marker status table was manually processed for areas with a high recombination frequency. These areas have been removed from the assembly. Markers belonging to deleted regions have also been removed from the marker state table.

At the next stage of data preparation for constructing a genetic map, the averages for markers within the contig were calculated for each contig and each sample. Based on the averages, a new table of marker states was compiled, in which contigs acted as markers. The following thresholds and a code corresponding to the input data of the MSTMAP software v. 1.0 (Wu *et al*., 2008) were used to transfer information about the average values of the states of markers in each specific position into the state of contig markers:

- “A” - homozygous allele corresponding to the reference. This state was assigned to a marker if the average for the contig corresponding to the marker in the sample was from -1 to -0.8, including the ends of the interval.
- “B” - homozygous allele corresponding to the homozygote from the second parent F0 plant of the genetic map. This state was assigned to a marker if the average for the contig corresponding to the marker in the sample was from 0.8 to 1, including the ends of the interval.
- “X” – heterozygote. This state was assigned to a marker if the average for the contig corresponding to the marker in the sample was from -0.2 to 0.2, including the ends of the interval.
- “-” – undefined marker. This state was assigned to a marker if the average for the contig corresponding to the marker in the sample was from -0.8 to -0.2 or from 0.2 to 0.8.

The resulting table of markers was processed in the MSTMAP software with the following parameters: “cut_off_p_value 0.0000001, no_map_dist 100.0, no_map_size 1, missing_threshold 0.5, population_type RIL2”. As a result of this program, a set of 15 linkage groups was obtained. For 5 linkage groups, the analysis was restarted by the MSTMAP software with the changed parameter “cut_off_p_value 0.0000000001”. As a result, 16 linkage groups were obtained.

To scaffold contigs using HiC, HiC reads were mapped onto a set of filtered contigs. Mapping was carried out without taking into account paired readings using the CLC program. For the obtained alignments, the information about the pairing of reads and information about the links was restored partly using the Arima mapping Pipeline (https://github.com/ArimaGenomics/mapping_pipeline Accessed 26 June 2023), partly using the scripts we developed. The obtained linkage information was further divided into linkage groups and separately processed using the AllHiC program (Zhang *et al*., 2019), which was modified to use information about the order of contigs in the linkage group. As a result, an assembly was obtained, consisting of 16 HiC scaffolds and contigs not included in them. Visualization of HiC maps was carried out using the JuiceBox program (Robinson *et al*., 2018). Based on the analysis of maps for each linkage group, scaffolding errors were identified and corrected.

### Genome annotation

Annotation was the multistep procedure that started from two annotations - one was done using BRAKER v. 2.1.5 (Hoff *et al*., 2019) with parameters “--esmode --softmasking” and the second - using liftoff v 1.6.1 (Shumate and Salzberg, 2021). Liftoff is a tool that can transfer annotations from closely related species (Arabidopsis thaliana, annotation TAIR10 in this case). The transfer was done separately for R and O subgenome, then the annotations for subgenomes were merged into one GFF file and all genes that did not contain valid ORF were removed using script RemoveNotValidORFsFromGffFile.pl. Then we added to this annotation the gene models generated with BRAKER using a script MergeGFFFiles.pl (only those genes models that did not overlap with filtered Liftoff annotation were added). For all genes annotated at this step CDS and corresponding amino acid sequences were predicted. The amino acid sequences were used for the inference of orthology with *A. thaliana* (TAIR10 proteins) using Orthofinder v 2.4.0, separately for each of the subgenomes. Then we considered orthogroups that contained one or several *C. bursa-pastoris*(Cbp) genes from either R or O subgenome and exactly one *A.thaliana* gene. This allowed to make a correspondence between Arabidopsis gene and Cbp genes; this correspondence was included in the gene name in order to facilitate further analyses. The genes that were not the part of the orthogroups with single A. thaliana genes (the genes from Cbp-only orthogroups, or orthogroups with more than one A. thaliana gene or singletones) were the subject of blastp search against TAIR10 proteins. The best hit was taken as a corresponding and its identifier was included in the Cbp gene name. At the nest step we searched for the pairs of homeologs. In order to identify them we first performed blastn search of all CDS from a chromosome (or part of the chromosome, in case of hybrid chromosomes R7_O7 and O7_R7) belonging to R subgenome against all CDS from homologous chromosome of the O subgenome and vice versa. Then using a script GetListOfSequenceGenesForGFFFileWithSequenceOfHomeologous.pl we compiled a list of pairs, using the information on the best blast hit and the position of a gene in the chromosome (i.e. if for example, a certain gene from R subgenome has two hits in the homologous chromosome of the O subgenome, the gene with the position that better corresponds to the position in the R subgenome is considered as a homeolog). For the gene for which the homeologs we not found we performed an additional round of gene prediction using liftoff. In this case liftoff transfers the annotation from a gene that was found but left without pair to a subgenome where the homeolog of this gene should be but was not found. For example, let’s consider the genes O1, O2 and O3 from the subgenome O; O1 and O3 have pairs of homeologs R1 and R3 correspondingly, and O2 does not have a homeolog. In this case we use liftoff to predict a gene in a region that is located between R1 and R3, giving it a hint that this gene should be highly similar to O2. After this, the genes that did not contain valid ORF were filtered using script RemoveNotValidORFsFromGffFile.pl and merged with the main annotation using script MergeGFFFiles.pl. Then the inference of orthology, BLAST search and the search of homeologs were run again with the set that includes these additionally discovered genes. Based on the information on homeologous relationships between R- and O subgenome genes and on the similarity with A. thaliana genes we named the genes in Cbp annotation using script ReformatGffFile.pl.

### Phylogenetic trees and networks

For the inference of relationships between *C. bursa-pastoris*, *C. rubella*/*C. grandiflora* and *C.orientalis* and within Cbp we mapped data from multiple individuals of these species on our assembly of *C. bursa-pastoris* genome. The data included sequences from earlier publications (Huang *et al*., 2018; Kryvokhyzha, Salcedo, *et al*., 2019) (and generated in this study. In order to provide reliable results of SNP calling we selected only those samples that had sequencing depth more than 10x and that represented individual plants, not pools of several neighboring plants. The data for *C.rubella*/*C. grandiflora* and *C.orientalis* were mapped separate on R subgenome and O subgenome, and for *C. bursa-pastoris* - on both subgenomes. For mapping and SNP calling we used the tool CLC Genomics workbench (Qiagen) (parameters identical described on Supplementary figure 7). Mapping and SNP calling setting were the same as used for the construction of genetic map (Supplementary figure 2). Then the lists of SNP were retrieved for each individuum (for Cbp samples - for each subgenome separately) using the script CreateDBList.pl. All SNPs found in at least one sample were added to the list. Then the sequencing depth was calculated for each SNP from the list using samtools depth. The SNPs with sequencing depth less than 4 were replaced by N; heterozygous SNPs were replaced by corresponding letters denoting degenerate bases (Y for C/T, R for A/G etc). These data were used for the construction of pseudoalignment. For each sample, the sequence was constructed based on the reference sequence with replacement of reference positions carrying SNP by the nucleotide corresponding to the non-reference nucleotide found in this sample. For this we used a custom script CreatePseudoAlignmentForRegion.pl. Separate pseudoalignments were constructed for each chromosome arm. Phylogenetic trees were constructed for each pseudoalignment using RAXML v. 8.0.26 with parameters “-m GTRCAT -T 100 -x 123456 -N 100 -p 092345”.

### Admixture analysis

VCF files with filtered SNPs obtained from the variant calling on the subgenome R and O of the *C. bursa-pastoris* “msk-wt” line reference genome were merged into two multisample VCF files for each subgenome separately using bcftools v.1.16 (Danecek *et al*., 2021) (*merge* with option *--missing-to-ref*). VCF files have contained samples of *C. rubella*, *C. grandiflora*, *C. orientalis*, and three major populations of *C. bursa-pastoris,* 102 samples in total in each file. Then VCFs were converted to plink binary format using plink software v1.90b6.21 (Chang, 2020; Chang *et al*., 2015) to infer the population structure using fastStructure v.1.0 software (Anon, n.d.), which accepts genotypes in plink binary format as input. Admixture proportions were calculated with or without parental species of *C. bursa-pastoris* with the K value (number of populations) from 1 to 10. The optimal model complexity was estimated using the chooseK Python script of the fastStructure software package. Admixture proportions inferred by fastStructure were visualized using distruct v.2.3 Python script (http://distruct2.popgen.org Accessed 26 June 2023).

## Data Availability Statement

Raw sequencing reads for are available in the NCBI under Bioproject PRJNA986448 (PacBio and HiC reads), PRJNA986297 (individual plants F2 lel x wt3.4-msk),PRJNA986442 (individual plants F2 wt3.4-msk x ME line). Genome assembly are available in https://figshare.com/s/82d547f4b5e180cfda1b. Custom scripts are available on github page https://github.com/ArtemKasianov/CapsellaArticle2

## Funding

This study was supported Russian Science Foundation, project # 21-74-20145.

## Supplementary figures

**Supplementary figure 1.** Scheme for data acquisition for the genetic map.

**Supplementary figure 2.** An example of a contig fragment with a colored state of the markers. Due to the low coverage some of the markers are "noisy".

**Supplementary figure 3.** An example of chimeric assembly. Two adjacent markers located at a distance of ∼90 kbp have 33 "recombinations" per 100 chromosomes, which is impossible and indicates independent inheritance of sites.

**Supplementary figure 4.** An example of correction of a local chimeric assembly - insertion of a foreign fragment(s) and view of the site after correction (b). The green dashed lines show the signals indicating the proximity of the sites.

**Supplementary figure 5.** Simulation of the subgenome separation procedure. Example of the coverage by reads of the parental species of some reference contigs created from the genomes of *C. orientalis* and *C. rubella* for subgenome separation.

**Supplementary figure 6.** Analysis of introgression by admixture analysis in *C. bursa-pastoris*, for K=6. The colors of the line names correspond to the populations in Figure 4b.

**Supplementary figure 7.** F2 data processing.

**Supplementary figure 8.** Building the SNP Database.

## Supplementary tables

**Supplementary table 1.** BUSCO metrics.

**Supplementary table 2.** Set of samples used for genetic variation analysis.

## Notes

https://github.com/ArtemKasianov/CapsellaArticle2

https://figshare.com/s/82d547f4b5e180cfda1b

### Competing Interest Statement

The authors have declared no competing interest.

