## Supplementary figures for "Origin and diversity of *Capsella bursa-pastoris* from the genomic point view"

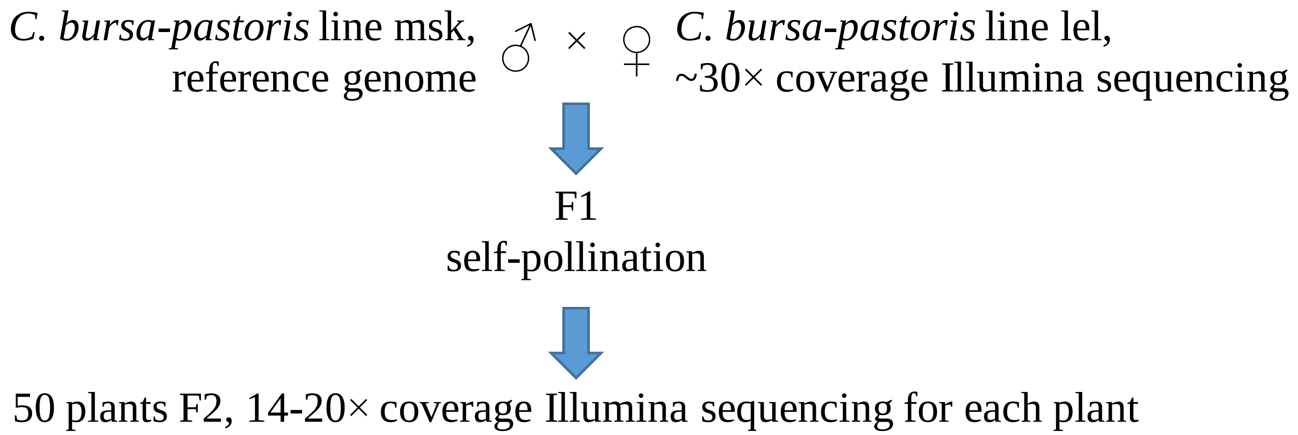


**Supplementary figure 1.** Scheme for data acquisition for the genetic map.

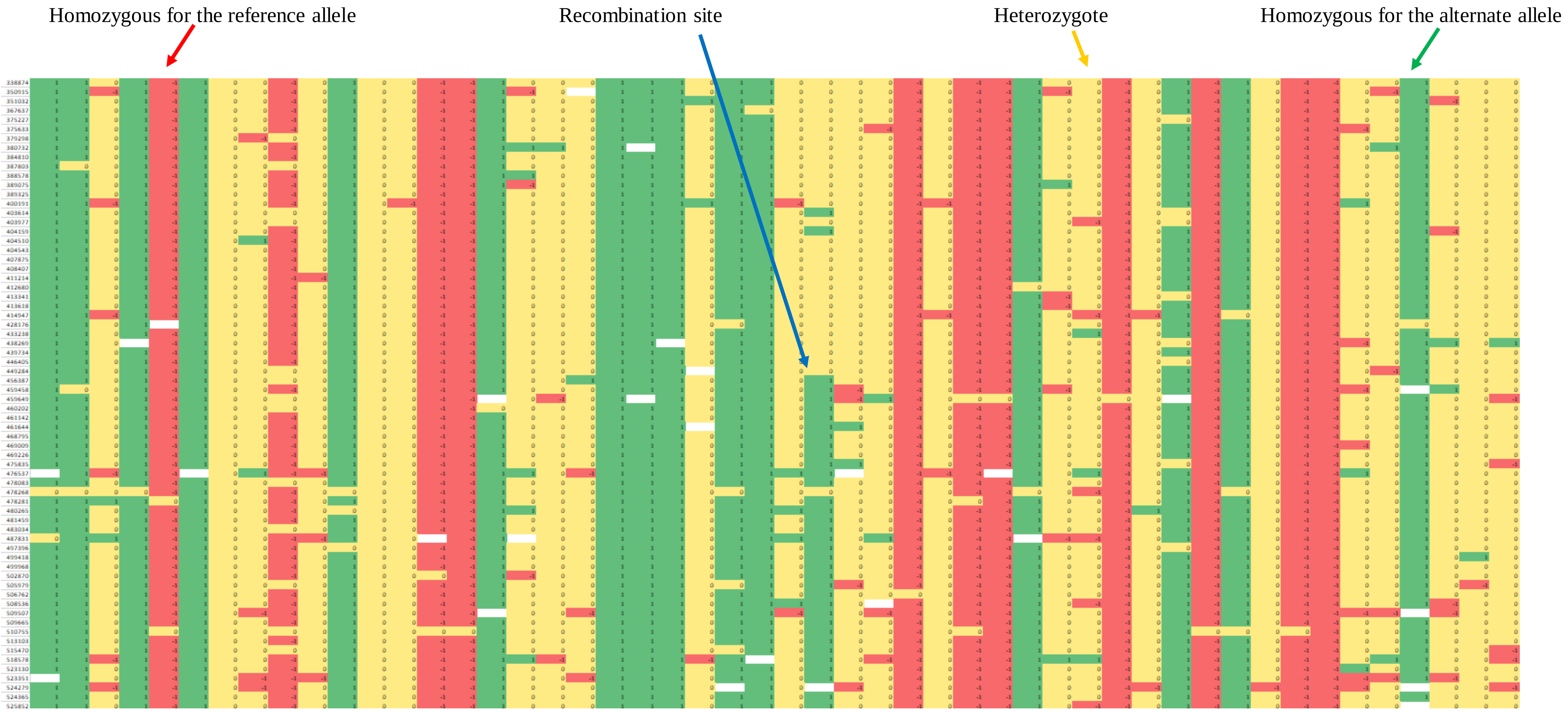


**Supplementary figure 2.** An example of a contig fragment with a colored state of the markers. Due to the low coverage some of the markers are "noisy".

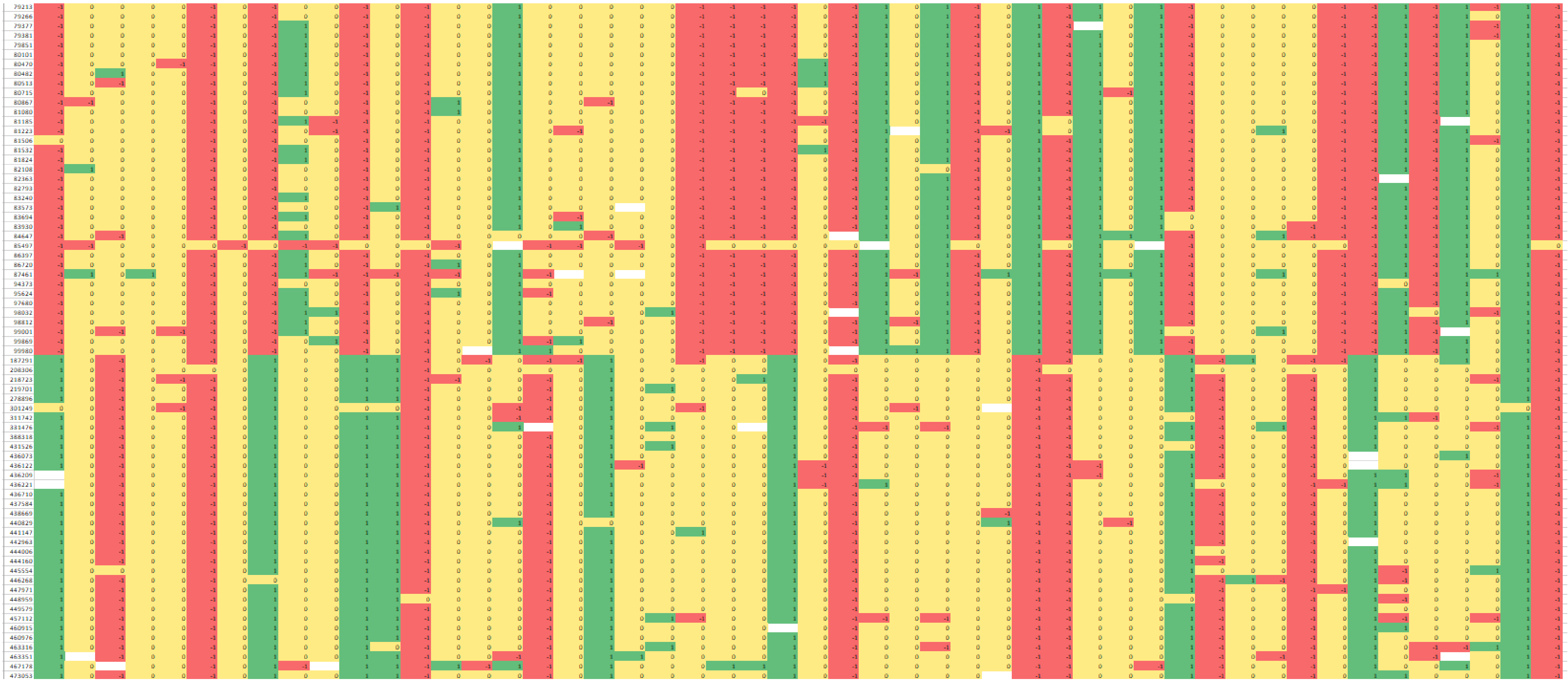

**Supplementary figure 3.** An example of chimeric assembly. Two adjacent markers located at a distance of ~90 kbp have 33 "recombinations" per 100 chromosomes, which is impossible and indicates independent inheritance of sites.

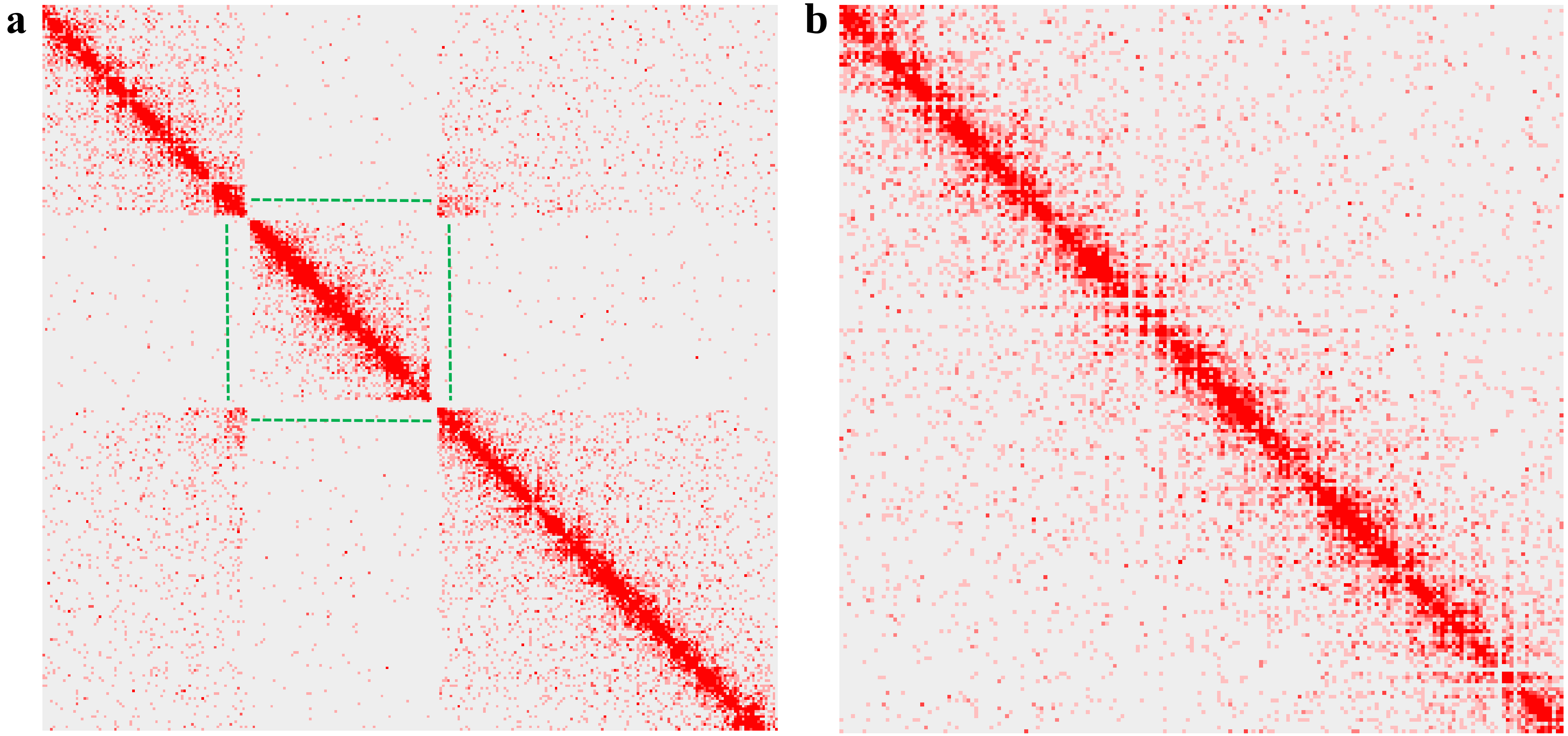

**Supplementary figure 4.** An example of correction of a local chimeric assembly - insertion of a foreign fragment(s) and view of the site after correction (b). The green dashed lines show the signals indicating the proximity of the sites.

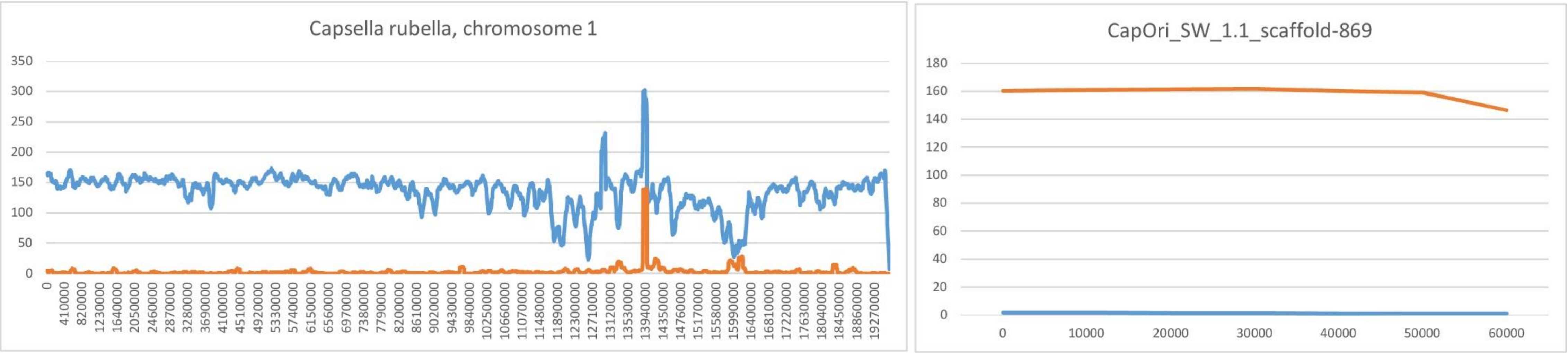


**Supplementary figure 5.** Simulation of the subgenome separation procedure. Example of the coverage by reads of the parental species of some reference contigs created from the genomes of C. orientalis and C. rubella for subgenome separation.

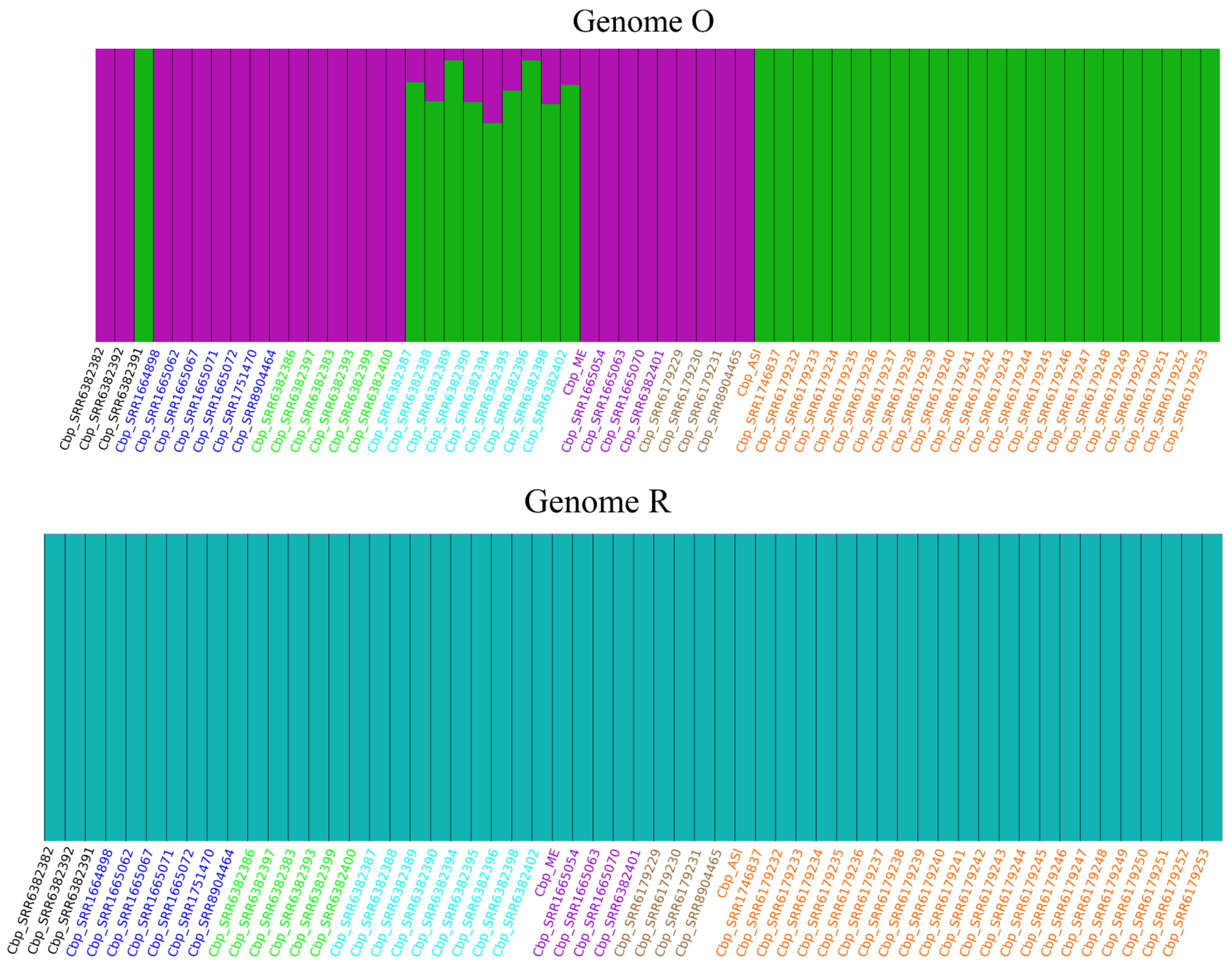


**Supplementary figure 6.** Analysis of introgression by admixture analysis in C. bursa-pastoris, for K=6. The colors of the line names correspond to the populations in Figure 4b.

### Mapping Illumina sequencing reads of the lel line to the reference genome using CLC Genomics Workbench 20.0.3

**Parameters**: Match score = 1, Mismatch cost = 3, Cost of insertions and deletions = Linear gap cost, Insertion cost = 3, Deletion cost = 3, Length fraction = 1.0, Similarity fraction = 0.93, Non-specific match handling = Ignore

### Alignment

Total mapped reads ~58-63%, in pairs ~51-58%,
>7× average coverage

### Basic variant detection 2.1. using CLC Genomics Workbench 20.0.3

**Parameters**: Ploidy = 2, Ignore positions with coverage above = 100, Ignore broken pairs = Yes, Ignore non-specific matches = Reads, Minimum coverage = 2, Minimum count = 4, Minimum frequency (%) = 15.0, Base quality filter = Yes, Neighborhood radius = 5, Minimum central quality = 20, Minimum neighborhood quality = 15 Read direction filter = Yes, Direction frequency (%) = 5.0, Relative read direction filter = Yes, Significance (%) = 1.0, Read position filter = Yes, Significance (%) = 1.0

### Filtration by the SNP database using following rules:

- All polymorphic sites with no matching record in the database were deleted.
- If the variant had been found in the database but not in the F2 plant, its coverage was checked. If the coverage was ≥4, then the reference allele was inserted, else the marker for this plant was set as unidentified.
- Only markers that were present in 90% of the F2 plants were taken.
- Only markers that passed the 1:1 ratio of alleles in F2 according to Pearson's criterion were taken.
- To view and analyze alleles were coded as "-1" for the reference, "1" for the alternative, "0" for heterozygote, and "-" for the markers with an unknown state.

**Supplementary figure 7.** F2 data processing.

Mapping Illumina sequencing reads of the lel line to the reference genome using CLC Genomics Workbench 20.0.3

**Parameters:** Match score = 1, Mismatch cost = 3, Cost of insertions and deletions = Linear gap cost, Insertion cost = 3, Deletion cost = 3, Length fraction = 1.0, Similarity fraction = 0.93, Non-specific match handling = Ignore

↓

Alignment

Total mapped reads 55.30%, in pairs 50.05%, 16× average coverage

↓

Basic variant detection 2.1. using CLC Genomics Workbench 20.0.3

**Parameters**: Ploidy = 2, Ignore positions with coverage above = 100, Ignore broken pairs = Yes, Ignore non-specific matches = Reads, Minimum coverage = 4, Minimum count = 4, Minimum frequency (%) = 100.0, Base quality filter = Yes, Neighborhood radius = 5, Minimum central quality = 20, Minimum neighborhood quality = 15, Read direction filter = Yes, Direction frequency (%) = 5.0, Relative read direction filter = Yes, Significance (%) = 1.0, Read position filter = Yes, Significance (%) = 1.0

↓

Total variants database

SNV - 254283, MNV - 3370, Insertion - 13252, Deletion – 16984, 12.2× average SNP coverage.

↓

SNP database

Use only SNV - 254283, Frequency ~0.82 on 1kbp

**Supplementary figure 8.** Building the SNP Database.
