## Supplementary tables for "Origin and diversity of *Capsella bursa-pastoris* from the genomic point view"

| The analyzed sequences | Complete BUSCOs viridiplantae (brassicales) | Complete and single-copy BUSCOs viridiplantae (brassicales) | Complete and duplicated BUSCOs viridiplantae (brassicales) | Fragmented BUSCOs viridiplantae (brassicales) | Missing BUSCOs viridiplantae (brassicales) | Total gene number |
|---|---|---|---|---|---|---|
| **Full assembly** | 98,6 (99,5) | 5,3 (5,0) | 93,3(94,5) | 0,2(0) | 1,2(0,5) | 65207 |
| **Chromosome** | 98,1 (99,4) | 7,9 (6,7) | 90,2(92,7) | 0,2(0,0) | 1,7(0,6) | 63776 |
| **Subgenome O** | 96,1 (97,2) | 93,3 (95,5) | 2,8(1,7) | 0,5(0,2) | 3,4(2,6) | 31875 |
| **Subgenome R** | 98,1 (97,8) | 95,8 (96,3) | 2,3(1,5) | 0,2(0,5) | 1,7(1,7) | 31901 |
| ***Arabidopsis thaliana*** | 99,0 (99.3) | 98,1 (98.0) | 0,9 (1,3) | 0 (0,1) | 1,0 (0,6) | 27636 |
| **Contigs excluded after correction using genetic map** | 2,1 (1,4) | 1,6 (1,2) | 0,5(0,2) | 0,5(0,1) | 97,4(98,5) | 659 |
| **Contigs not included in linkage groups** | 0,2 (0) | 0,2(0) | 0(0) | 0(0) | 99,8(100) | 4 |
| **Sequences remoted after of HiC correction** | 2,3 (2,1) | 2,1(2,1) | 0,2(0) | 0,2(0) | 97,5(97,9) | 768 |
| **Sequences in total not included in pseudochromosomes** | 4,7 (3,6) | 4(3,3) | 0,7(0,3) | 0,2(0,1) | 95,1(96,3) | 1431 |

**Supplementary table 1.** BUSCO metrics

| SRA | BioSample | Population | Geographic location | Latitude | Longitude | Group number |
|---|---|---|---|---|---|---|
| **SRR6382390** | SAMN08193318 | ASI | China | 26,37 | 106,43 | Group6 |
| **SRR6382398** | SAMN08193330 | ASI | China | 33,57 | 107,45 | Group6 |
| **SRR6382389** | SAMN08193319 | ASI | China | 30,16 | 120,13 | Group6 |
| **SRR6382391** | SAMN08193321 | ASI | China | 26,53 | 112,33 | Group7 |
| **SRR1665070** | SAMN03225186 | ME | Italy: Bacia | 43,00 | 12,55 | Group3 |
| **SRR1665063** | SAMN03225183 | ME | Greece: Artemida | 37,97 | 24,00 | Group3 |
| **SRR1665054** | SAMN03225184 | ME | Spain: Valladolid | 41,69 | -4,73 | Group3 |
| **SRR6382401** | SAMN08193333 | ME | USA | 31,29 | -97,17 | Group3 |
| **CBP_ME** | - | ME | UK: London | 51,48 | -0,29 | Group3 |
| **SRR6382386** | SAMN08193314 | ME | Algeria | 35,45 | 7,96 | Group2 |
| **SRR6382397** | SAMN08193331 | ME | Turkey | 41,02 | 28,97 | Group2 |
| **SRR6382383** | SAMN08193322 | EU | Russia | 52,16 | 104,18 | Group2 |
| **SRR6382393** | SAMN08193327 | EU | Sweden | 56,15 | 13,77 | Group2 |
| **SRR6382399** | SAMN08193329 | EU | France | 44,51 | -1,21 | Group2 |
| **SRR6382400** | SAMN08193328 | EU | United Kingdom | 56,20 | 2,47 | Group2 |
| **SRR6179229** | SAMN07792208 | EU | China: Buerjin | 47,70 | 86,85 | Group4 |
| **SRR6179230** | SAMN07792207 | EU | China: Fuyun | 47,00 | 89,53 | Group4 |
| **SRR6179231** | SAMN07792206 | EU | China: Tacheng | 46,78 | 82,98 | Group4 |
| **SRR8904465** | SAMN11417657 | EU | China | 47,07 | 83,01 | Group4 |
| **SRR1665071** | SAMN03225189 | EU | Russia: Vladivostok | 43,13 | 131,91 | Group1 |
| **SRR1665067** | SAMN03225185 | EU | Poland: Krakow | 50,06 | 19,95 | Group1 |
| **SRR1751470** | SAMN03280713 | EU | Sweden: Harnasand | 62,65 | 17,89 | Group1 |
| **SRR8904464** | SAMN11417656 | EU | Russia | 66,65 | 66,40 | Group1 |
| **SRR1665072** | SAMN03225187 | EU | Germany: Halle | 51,49 | 11,98 | Group1 |
| **SRR1665062** | SAMN03225188 | EU | Netherlands: Nijmegen | 51,81 | 5,85 | Group1 |
| **SRR1664898** | SAMN03225194 | EU | Iceland: Reykjavik | 64,15 | -21,94 | Group1 |
| **SRR6382382** | SAMN08193317 | EU | France | 48,08 | 7,37 | NA |
| **SRR6382392** | SAMN08193320 | EU | China | 45,45 | 126,37 | NA |

**Supplementary table 2.** Set of samples used for genetic variation analysis, page 1

| SRA | BioSample | Population | Geographic location | Latitude | Longitude | Group number |
|---|---|---|---|---|---|---|
| SRR1746837 | SAMN03225190 | ASI | Taiwan: Puli | 24,00 | 120,96 | Group5 |
| SRR6179252 | SAMN07792184 | ASI | China: Fuzhou | 26,07 | 119,30 | Group5 |
| SRR6179238 | SAMN07792196 | ASI | China: Wuhan | 30,59 | 114,30 | Group5 |
| SRR6179244 | SAMN07792192 | ASI | China: Jiujiang | 29,66 | 115,95 | Group5 |
| SRR6179237 | SAMN07792199 | ASI | China: Nanjing | 32,10 | 118,79 | Group5 |
| SRR6179240 | SAMN07792194 | ASI | China: Huangshi | 30,20 | 115,03 | Group5 |
| SRR6179239 | SAMN07792197 | ASI | China: Shanghai | 31,24 | 121,48 | Group5 |
| SRR6179236 | SAMN07792198 | ASI | China: Hefei | 31,82 | 117,23 | Group5 |
| SRR6179241 | SAMN07792195 | ASI | China: Anqing | 30,55 | 117,06 | Group5 |
| SRR6179243 | SAMN07792203 | ASI | China: Changbaishan | 42,03 | 128,07 | Group5 |
| SRR6179233 | SAMN07792204 | ASI | China: Haerbin | 45,80 | 126,63 | Group5 |
| SRR6179232 | SAMN07792205 | ASI | China: Haerbin | 45,80 | 126,63 | Group5 |
| SRR6179234 | SAMN07792200 | ASI | China: Qingdao | 36,11 | 120,38 | Group5 |
| SRR6179235 | SAMN07792201 | ASI | China: Handan | 36,63 | 114,54 | Group5 |
| SRR6179242 | SAMN07792202 | ASI | China: Huangyuan | 36,68 | 101,25 | Group5 |
| SRR6179245 | SAMN07792193 | ASI | China: Lulang | 29,94 | 94,80 | Group5 |
| SRR6179247 | SAMN07792191 | ASI | China: Baimaxueshan | 28,37 | 99,02 | Group5 |
| SRR6179246 | SAMN07792190 | ASI | China: Zhongdian | 27,84 | 99,74 | Group5 |
| SRR6179249 | SAMN07792189 | ASI | China: Zhongdian | 27,84 | 99,74 | Group5 |
| SRR6179248 | SAMN07792188 | ASI | China: Lijiang | 26,86 | 100,22 | Group5 |
| SRR6179253 | SAMN07792185 | ASI | China: Huize | 26,42 | 103,30 | Group5 |
| SRR6179250 | SAMN07792186 | ASI | China: Zhijin | 26,66 | 105,78 | Group5 |
| SRR6179251 | SAMN07792187 | ASI | China: Guiyang | 26,65 | 106,64 | Group5 |
| CBP_ASI | NA | ASI | China: Kunming | 25,14 | 102,74 | Group5 |
| SRR6382396 | SAMN08193324 | ASI | China | 30,20 | 112,06 | Group6 |
| SRR6382402 | SAMN08193332 | ASI | China | 43,13 | 131,40 | Group6 |
| SRR6382388 | SAMN08193316 | ASI | China | 38,56 | 121,35 | Group6 |
| SRR6382387 | SAMN08193334 | ASI | China | 36,37 | 101,46 | Group6 |
| SRR6382395 | SAMN08193325 | ASI | China | 25,06 | 102,41 | Group6 |
| SRR6382394 | SAMN08193326 | ASI | China | 32,03 | 118,46 | Group6 |

**Supplementary table 2.** Set of samples used for genetic variation analysis, page 2
